# Unlocked capacity of proteins to attack membranes characteristic of aggregation: the evil for diseases and aging from Pandora’s box

**DOI:** 10.1101/071274

**Authors:** Liangzhong Lim, Yimei Lu, Jianxing Song

## Abstract

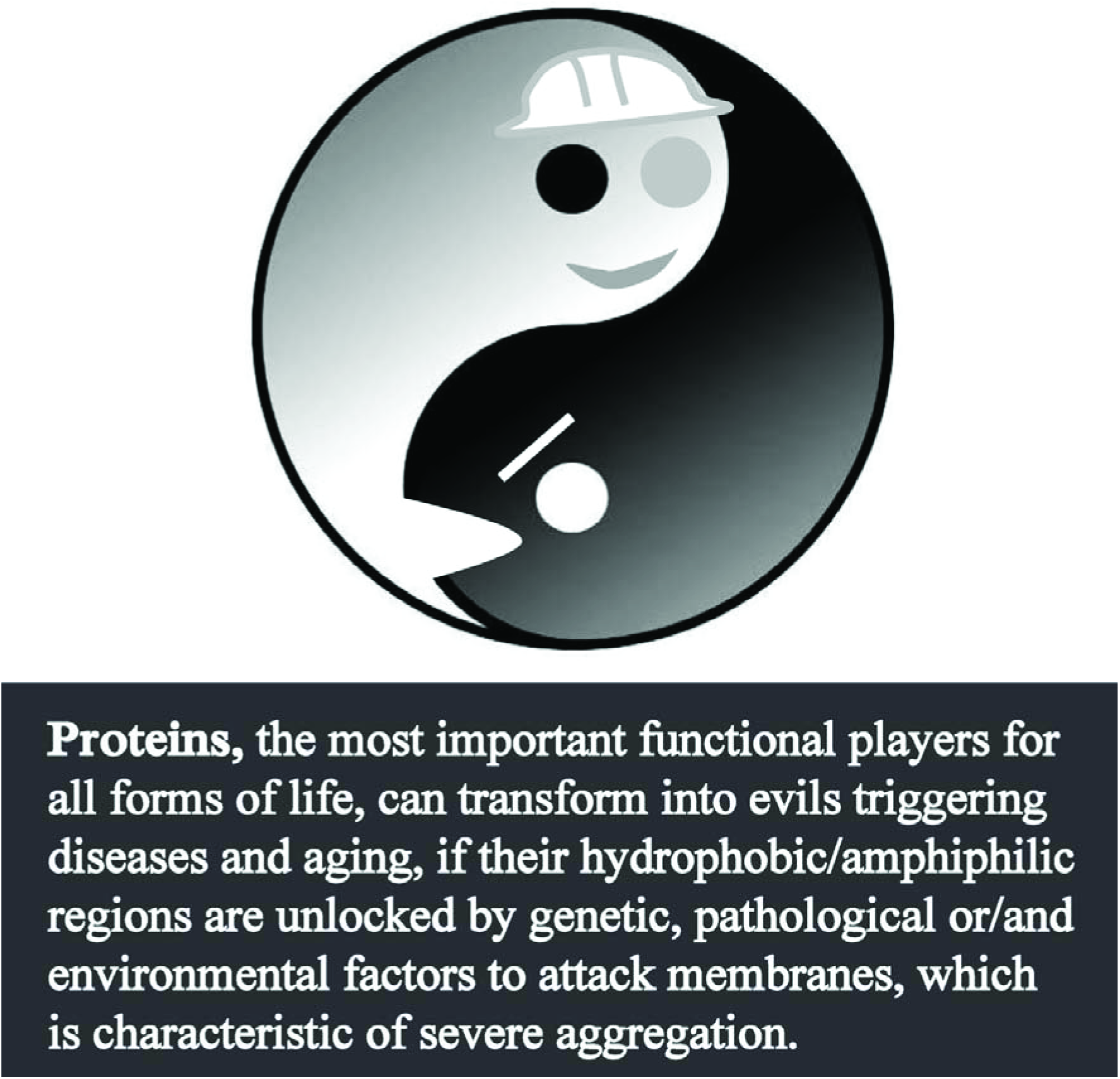

**Abstract:** Aggregation of specific proteins is characteristic of a large spectrum of human diseases including all neurodegenerative diseases, while aggregation of non-specific proteins has been now identified to be a biomarker for cellular aging down to *Escherichia coli*. Previously, as facilitated with our discovery in 2005 that “completely insoluble” proteins could be all solubilized in unsalted water [Song (2009) *FEBS Lett*. 583: 953], we found that the TDP-43 prion-like domain in fact contains an intrinsic membrane-interacting subdomain [Lim et al. [2016] *PLoS Biol.* 14, e1002338]. We decrypted that ALS-causing mutations/cofactor-depletion act to render the well-structured folds of cytosolic VAPB-MSP domain and SOD1 into highly disordered states, thus becoming buffer-insoluble. Most surprisingly, this also unlocks the amphiphilic/hydrophobic regions universally exiting in proteins, which thus acquire a novel capacity in abnormally interacting with membranes [Qin et al. (2013) *F1000Res* 2-221.v2; Lim (2016) *BBA-Biomembranes.* 1858: 2223]. Here we aimed extend our discovery to address two fundamental questions: 1) why many *E. coli* proteins become aggregated in aging; and 2) whether aggregation-prone proteins can also acquire a novel capacity in interacting with membranes; by dissecting the 557-residue S1 ribosomal protein into 7 fragments to disrupt its 6 S1 folds, followed by extensive CD and NMR characterizations. The results reveal that we have successfully eliminated all 6 S1 folds and fragment 4 becomes highly disordered and thus buffer-insoluble. Most strikingly, F4 does acquire a capacity in transforming into a helical conformation in membrane environments. Here, for the first time, our study deciphers that like ALS-causing mutants, the disruption of a well-folded *E. coli* cytosolic protein also unlocks its amphiphilic/hydrophobic regions which are capable of abnormally interacting with membranes. Therefore, proteins, the most important functional players for all forms of life, can transform into membrane-toxic forms triggering diseases and aging, if their hydrophobic/amphiphilic regions are unlocked by genetic, pathological or/and environmental factors, which is characteristic of severe aggregation.

## Introduction

Proteins are the most important functional players for all forms of life we know of so far. They are linear heteropolymers composed of 20 common α-amino acids, which amazingly are all in L-mirror-image [1]. Although 20 amino acids have distinctive characteristics, they can be briefly divided into two groups: hydrophobic (or non-polar) and hydrophilic (or polar). A portion of proteins spontaneously self-organizes into unique threedimensional structures via protein folding processes [2-4], while many are fully functional, but lack well-defined structures, and are thus called intrinsically disordered proteins (IDPs) [5]. It is widely recognized that the folding of cytosolic proteins is mainly resulting from solvophobic interactions of polar water molecules with the hydrophobic side chains of proteins [3,4].

One intriguing phenomenon associated with proteins is their insolubility in aqueous buffers. Protein aggregation/insolubility is not only problematic for *in vitro* protein research and industry applications, but is commonly characteristic of a large spectrum of human diseases [7-14], which include all human neurodegenerative diseases [7-11], such as Parkinson’s disease (PD), Alzheimer’s disease (AD), Huntington’s disease (HD), spinocerebellar ataxias (SCA), amyotrophic lateral sclerosis (ALS); as well as diabetes [12] and cardiac dysfunction [14]. Amazingly, aggregation of non-specific proteins have been now identified to be associated with aging of all organisms [15,16]; and in particular the cellular aging and rejuvenation of ***Escherichia coli*** cells has been found to be characteristic of asymmetric segregation of protein aggregates [17].

It has been widely established that aggregation of specific proteins leads to human diseases by “loss of functions” or/and “gain of functions”. However, as even for the human neurodegenerative diseases, the involved proteins are so functionally diverse and their knockout (such as SOD1) does not always result in the corresponding diseases. Consequently it is most likely there is no common mechanism underlying “loss of functions”. On the other hand, as proteins involved in the diseases are all prone to aggregation, it was widely thought that the formation of protein aggregates represents a common mechanism to trigger the diseases. However, some emerging evidence do not support this hypothesis as: 1) many neurodegenerative diseases such as amyotrophic lateral sclerosis (ALS) triggered by SOD1 mutants were initiated without any detectable protein aggregates [6,9,18]. 2) More radically, it has been increasingly found that inclusion body formation in fact reduces neuronal deaths [19,20].

Previously, partly-soluble proteins involved in human diseases have been found to have capacity to interact with membranes [8,12,18,21-30]. However, many disease-causing mutants were found to be “completely insoluble”, thus could not be studied before. However, in 2005, we discovered that all insoluble proteins, even the most hydrophobic integral membrane protein fragment in nature, are all soluble in pure water at concentrations at least up to 100 μM [31-52]. With this powerful tool, we have been recently focused on characterizing “completely insoluble” mutants/proteins causing ALS [40-51]. We deciphered that the TDP-43 prion-like domain in fact contains a hidden region which is capable of interacting with membranes [50]. We also decoded that ALS-causing mutation, or truncation, or cofactor-depletion acts to disrupt the well-structured native β-folds and consequently they become highly disordered in unsalted water but rapidly aggregated in buffers. Most surprisingly, the mutation/truncation/cofactor-depletion unlocks the amphiphilic/hydrophobic regions universally existing in all proteins, which thus acquire a novel capacity in abnormally interacting with membranes, although these proteins are all well-folded cytosolic proteins and their native functions have nothing to do with membrane interactions [43-45,52].

In the present study, we aimed to extend our discovery to address a fundamental question: whether aggregation of non-specific proteins associated with *Escherichia coli* aging also lead to unlocking the amphiphilic/hydrophobic regions of these proteins, thus acquiring a novel capacity to abnormally interact with membranes. To achieve this, we have examined the aggregation of proteins associated with cellular aging of *E. coli* cells as asymmetric segregation of protein aggregates is the only biomarker characteristic of its cellular aging and rejuvenation [17]. We decided to select a protein for detailed studies which satisfies three criteria: 1) the protein should be extensively found in the *E. coli* aggregation lists. 2) It contains well-structured folds rich in β-sheets. 3) It owns no membrane-interacting region and its native functions have nothing to do with membrane-interaction. As such, the S1 ribosomal protein was selected as it was identified to aggregate severely in an *E. coli* strain carrying the mutant of YajL protein [53], a prokaryotic homolog of parkinsonism-associated protein DJ-1 which functions to repair proteins in response to the global stress particular oxidative stress with multi-functions such as acting as a covalent chaperone [53-56]. Furthermore, the aggregate of S1 ribosomal protein was even identified in healthy *E. coli* cells [57].

As shown in Fig 1, S1 ribosomal protein contains 6 well-structured S1 domains linked by flexible regions [57,58]. To mimic fragmentation resulting from radical-mediated protein oxidation associated with aging [60-63], we dissected S1 ribosomal protein into 7 fragments with the cleavage sites all located within the well-structured S1 folds (Fig 1), in an attempt to eliminate the folded structures of all 6 S1 domains. We obtained recombinant proteins of the full-length S1 ribosomal protein and its 7 dissected fragments, which were then extensively characterized by CD and NMR spectroscopy as we previously conducted on other proteins [30-52,64,65]. The results reveal that the successful elimination of the S1 folds indeed leads to unlocking the amphiphilic/hydrophobic regions which acquire a novel capacity in abnormally interacting with membranes. Therefore, proteins, the most important functional players for all forms of life, can transform into membrane-toxic forms triggering diseases and aging, if their hydrophobic/amphiphilic regions are unlocked by genetic, pathological or/and environmental factors to attack membranes, which is characteristic of severe aggregation.

**Fig 1.**
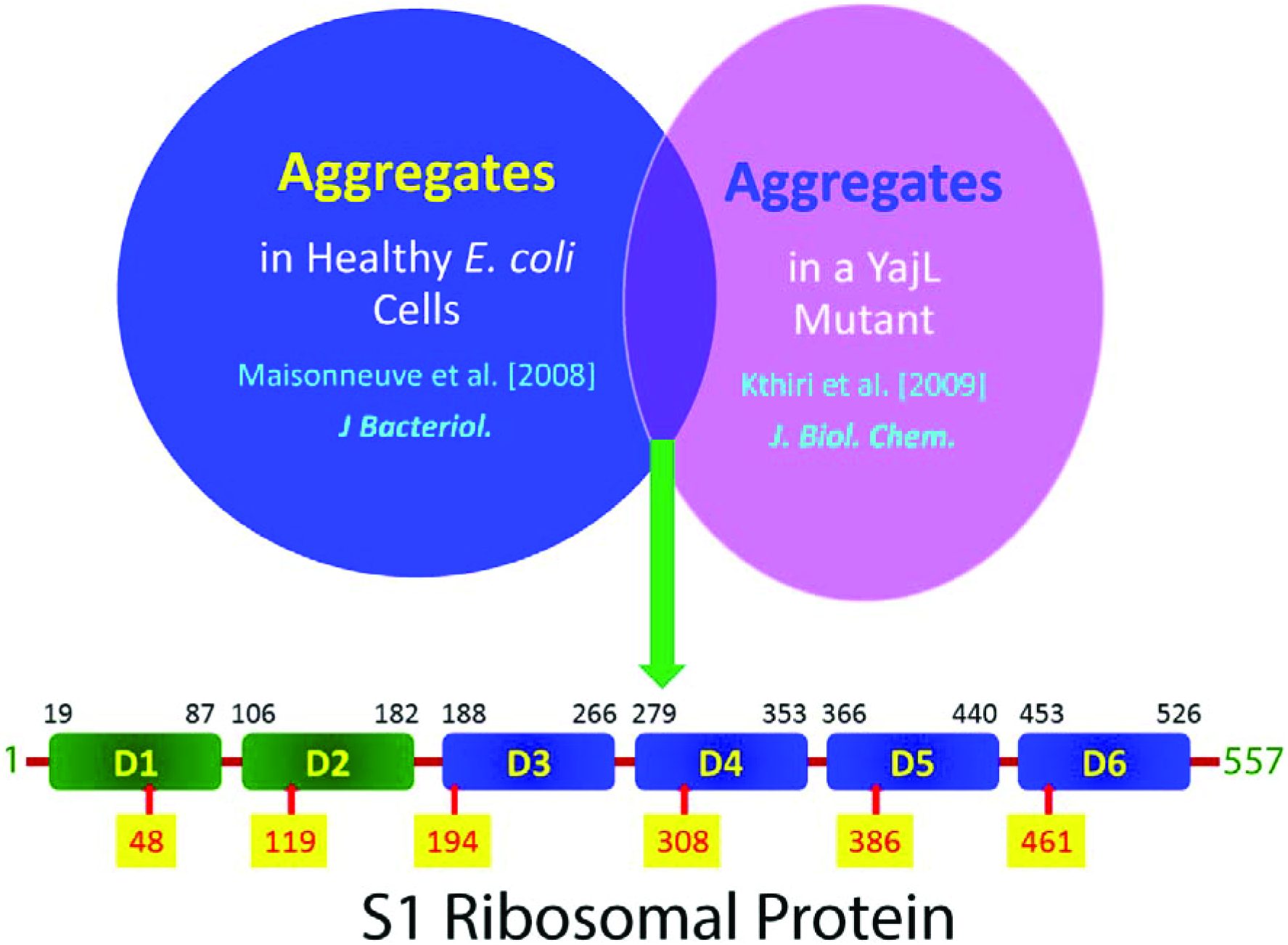
Selection and domain organization of *E. coli* S1 ribosomal protein. The S1 ribosomal protein was found in lists of protein aggregates of both healthy *E. coli* cells and of a YajL mutant cell strain. *E. coli* S1 ribosomal protein consists of 6 well-structured S1 domains. The starting residues for dissection are labeled, which were all located within the 6 S1 domains.

## Results

### 1. Dissection successfully eliminated the folded structures of six S1 domains

To mimic the fragmentation due to protein damages such as by radical-mediated overoxidation, we dissected the *E. coli* S1 ribosomal protein into 7 fragments with all 6 cleavage sites located within 6 S1 domains (Fig 1), which aimed to abolish the well-structured S1 fold. Subsequently we have successfully expressed and purified recombinant proteins of the fulllength and its 7 dissected fragments. Interestingly, except for the forth fragment (F4) which was only soluble in Milli-Q water, all the rest could be first dissolved in Milli-Q water and subsequently diluted into 1 mM phosphate buffer at pH 6.8 to reach a concentration of at least 100 μM. Fig 2 presents far-UV CD spectra of the full-length and 7 fragments at a concentration of 15 μM in both Milli-Q water (pH 4.0) and 1 mM phosphate buffer (pH 6.8).

**Fig 2.**
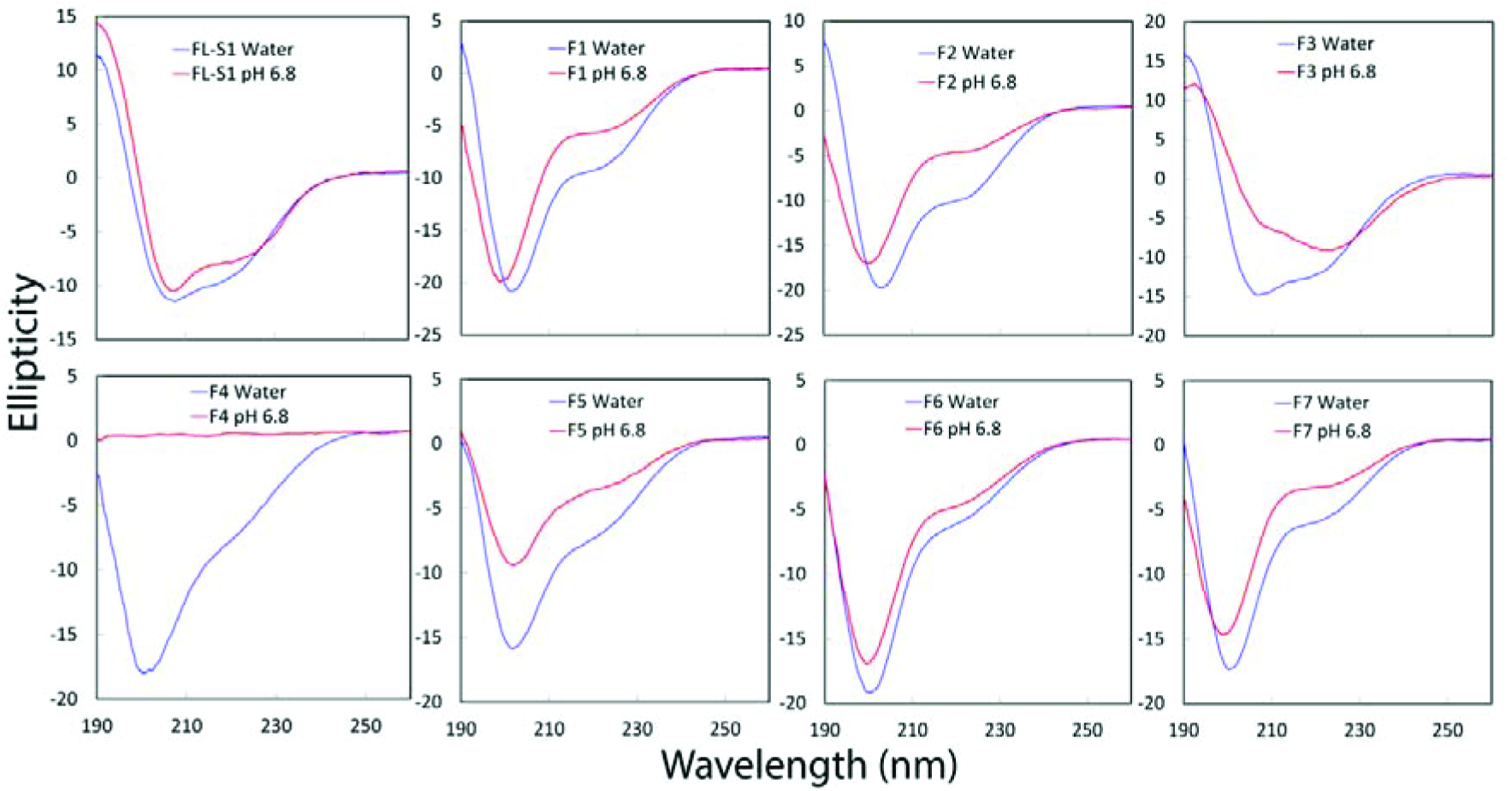
CD characterization. Far-UV CD spectra of the full-length S1 protein and its 7 dissected fragments in Milli-Q water (blue) and in 1 mM phosphate buffer at pH 6.8 (red).

However, F4 became completely precipitated upon dilution into the buffer and consequently no CD signal was detected in the buffer.

We further utilized NMR HSQC spectroscopy to characterize the solution conformations of the full-length and 7 fragments at a concentration of 100 μM in both Milli-Q water (pH 4.0) and 1 mM phosphate buffer (pH 6.8). As shown in Fig 3, probably due to the μs-ms conformational exchanges or/and dynamic aggregation, even in Mill-Q water the full-length has an HSQC spectrum lacking of well-dispersed NMR peaks typical of a well-folded proteins. In the buffer, most NMR peaks became too broadened to be detected (Fig 3), implying that salt ions can significantly enhance μs-ms conformational exchanges or/and dynamic aggregation as we previously deciphered on other proteins [31-52], although the secondary structures remain similar in Milli-Q water and the buffer as reflected by CD spectra (Fig 2). Most strikingly, all 7 dissected fragments had HSQC spectra which are also absent of any well-dispersed NMR peaks which were previously observed on isolated S1 domains [57,58]. This clearly indicates that our dissection successfully eliminated the ability of all 7 fragments to fold into the native S1 structures.

**Fig 3.**
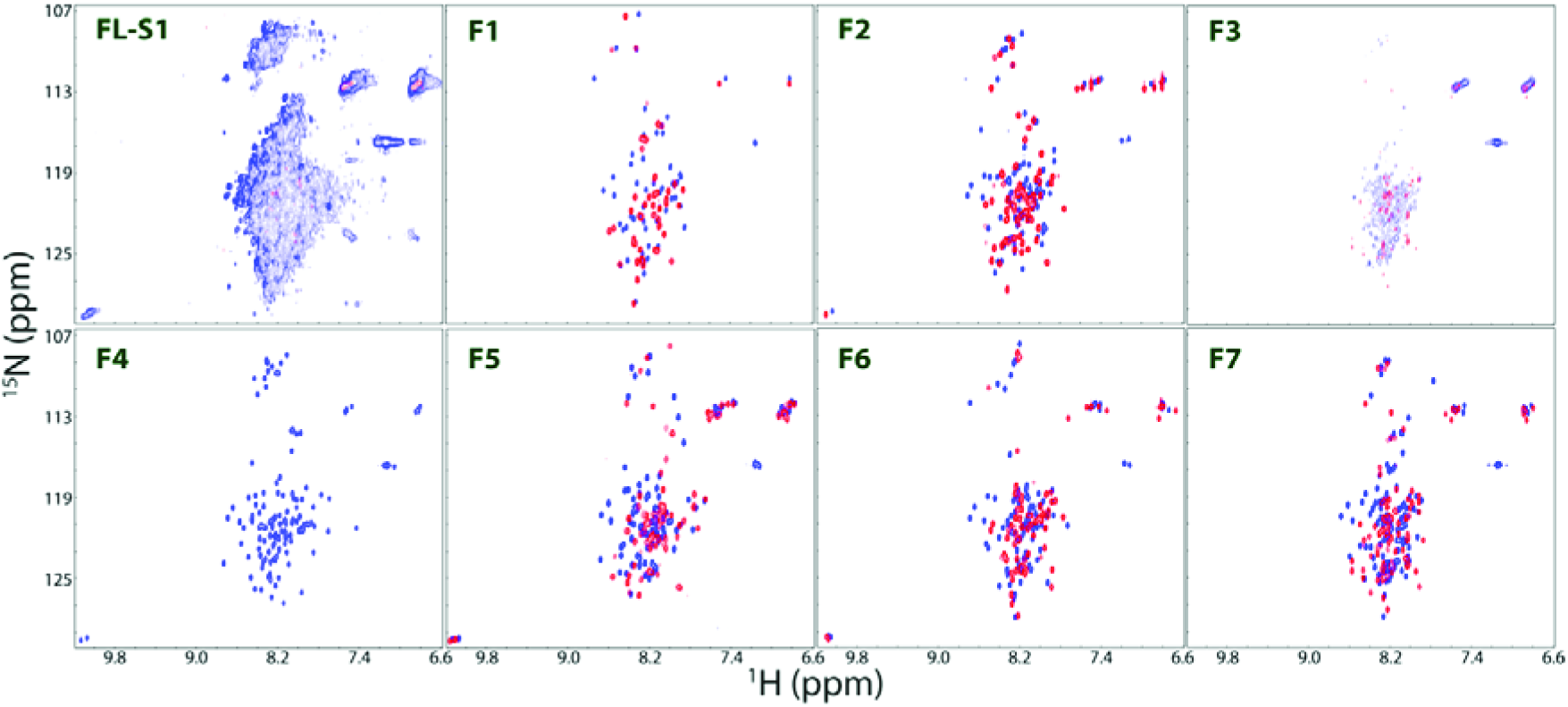
NMR HSQC characterization. NMR HSQC spectra of the full-length S1 protein and its 7 dissected fragments in Milli-Q water (blue) and in 1 mM phosphate buffer at pH 6.8 (red).

### 2. Interactions with DMPC/DHPC bicelle and liposome

To assess whether the full-length and 7 fragments have potential to interact with membranes, we titrated them with DMPC/DHPC bicelle as monitored by CD spectroscopy as we previously conducted on ALS-causing P56S-MSP mutant and TDP-43 prion-like domain. As shown in Fig 4, the full-length only had very small changes of the CD spectra upon adding the bicelle at different ratios. Out of 7 fragments, only F4 and F5 showed significant changes of the CD spectra upon adding the bicelle. In particular, as jud ed by CD spectra, F4 had a significant conformational transition from a highly disordered state in aqueous solution to helical conformation in bicelle. Furthermore, F4 also underwent a similar conformational transition upon interacting with liposome prepared from the total extract of *E. coli* lipids.

**Fig 4.**
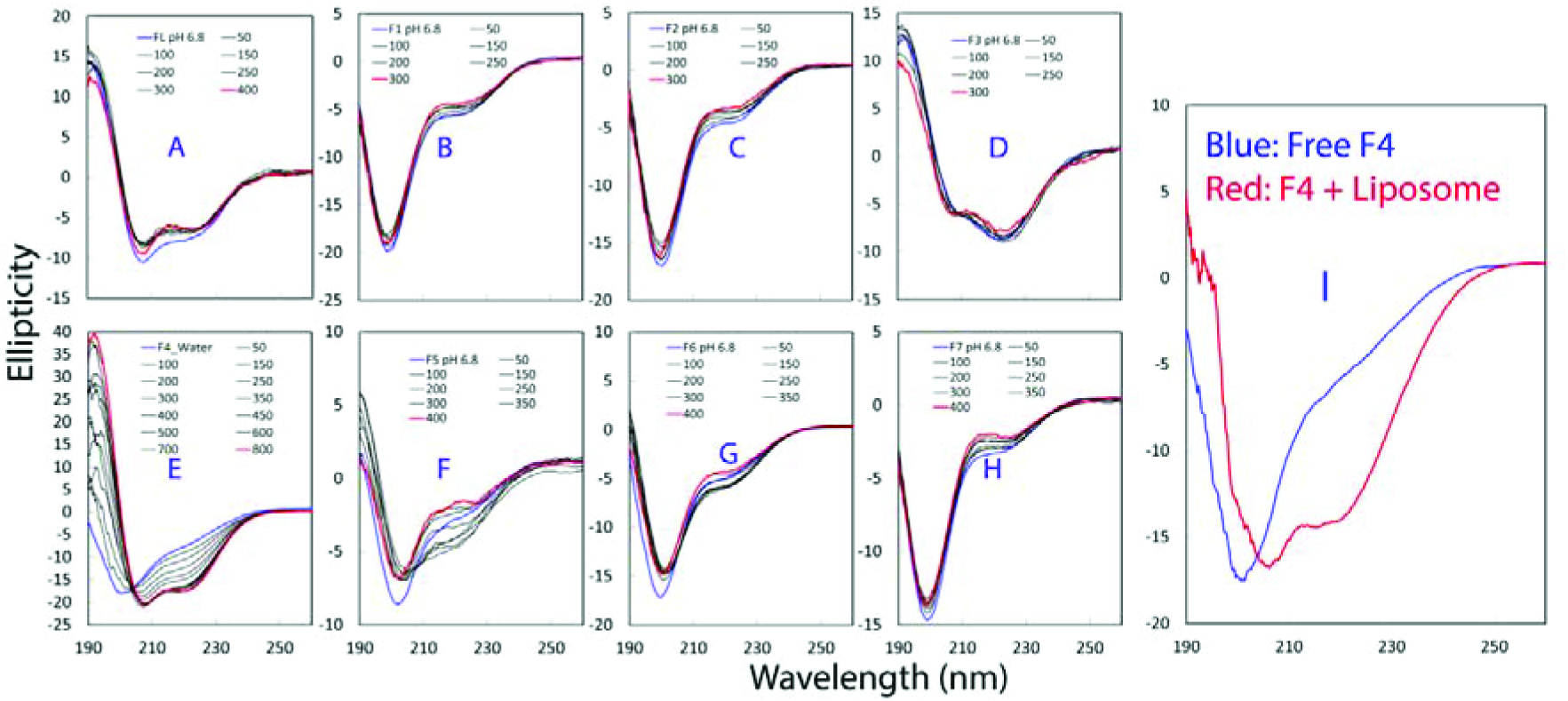
CD characterization of interactions with bicelle and liposome. (A-H) Far-UV CD spectra of the full-length S1 protein and its 7 dissected fragments in the presence of DMPC/DHPC bicelle at different molar ratios. (I) Far-UV CD spectra of F4 in the absence (blue) and in the presence (red) of liposome prepared from the total extract of *E coli* lipids.

As F4 was found to become rapidly precipitated in buffers, as well as to undergo significant conformational changes upon interacting with bicelle, we decided to further characterize its interaction with bicelle by NMR HSQC spectroscopy. As seen in Fig 5, HSQC peaks of F4 shifted significantly upon adding bicelle and the shift became largely saturated at a molar ratio of 1:200 (F4:bicelle). This clearly reveals that F4 was able to interact with bicelle to undergo a significant conformational change, consistent with the CD results (Fig 4).

**Fig 5.**
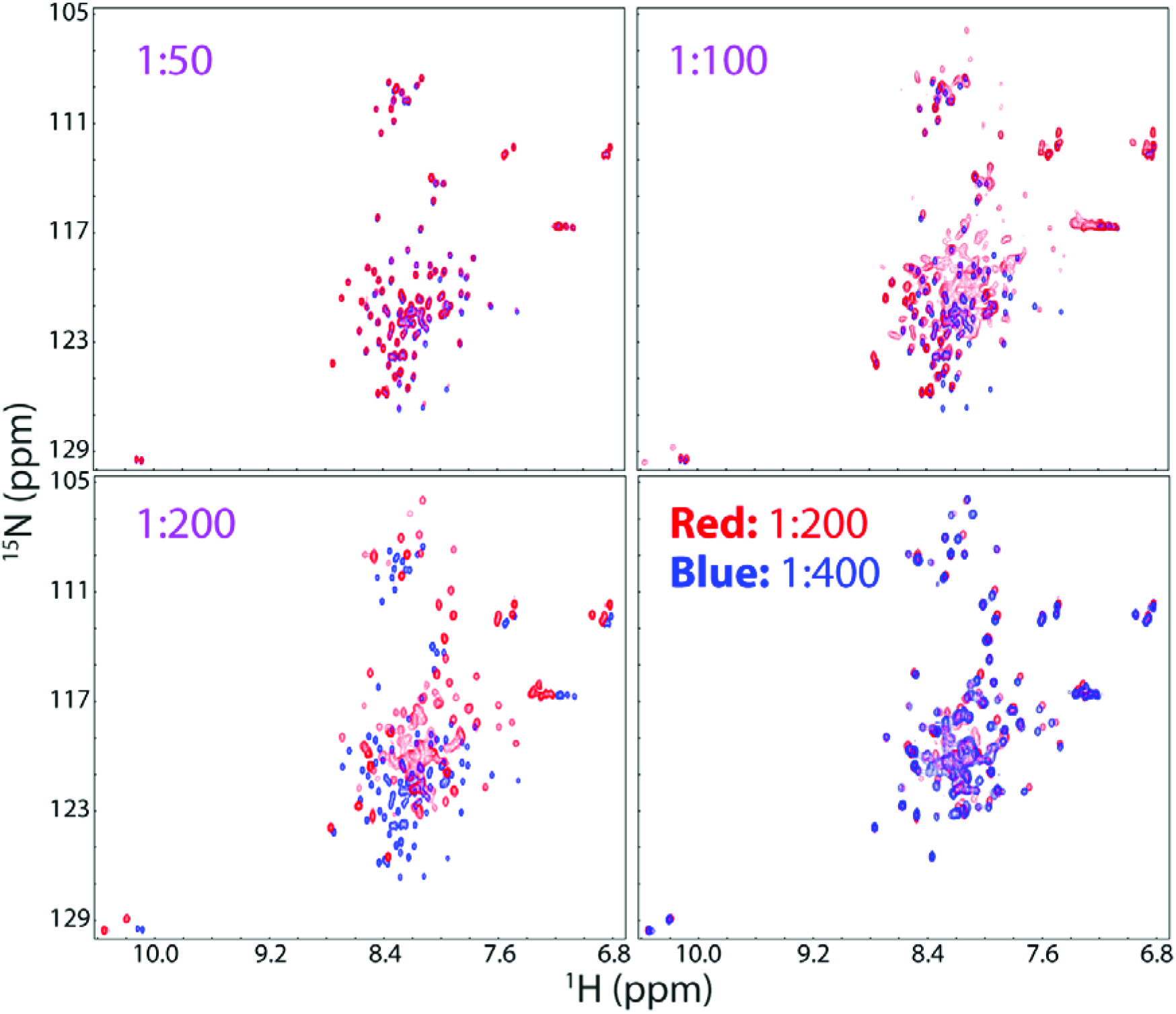
NMR HSQC characterization of interactions of F4 with Bicelle. NMR HSQC spectra of the fragment 4 in the presence of DMPC/DHPC bicelle at different molar ratios.

### 3. Residue-specific conformations of F4 in aqueous solution and bicelle

To gain insights into the conformational changes of F4 upon interacting with bicelle, by analyzing three-dimensional NMR spectra including CCC(CO)NH, HN(CO)CACB, here we successfully achieved NMR sequential assignments of F4 in aqueous solution and in bicelle. Fig 6 presents their (ΔCα-ΔCβ) chemical shifts, which represent a sensitive indicator of the residual secondary structures in disordered proteins [66]. In aqueous solution, F4 has small absolute values of (ΔCα-ΔCβ) chemical shifts over the whole sequence (Fig 6A), which clearly indicates that it is indeed lacking of any stable secondary structure, completely consistent with the CD results (Fig 4). To gain quantitative insights into the populations of different secondary structures, we further analyzed its NH, N, Hα, Cα and Cβ chemical shifts by SSP program [67]. As seen in Fig 6B, all residues have the absolute values of SSP scores less than 0.4, confirming that the whole domain has no stable secondary structure. Interestingly, the C-terminal residues over 279-306 of F4 adopt three β-strands in the forth S4 domain as previously determined by NMR spectroscopy [58,59]. This suggests that the abolishment of the well-structured folds leads to a complete loss of the β-stranded secondary structures, as we extensively observed on SH3, P56S-MSP and SOD1 mutants [31-52].

**Fig 6.**
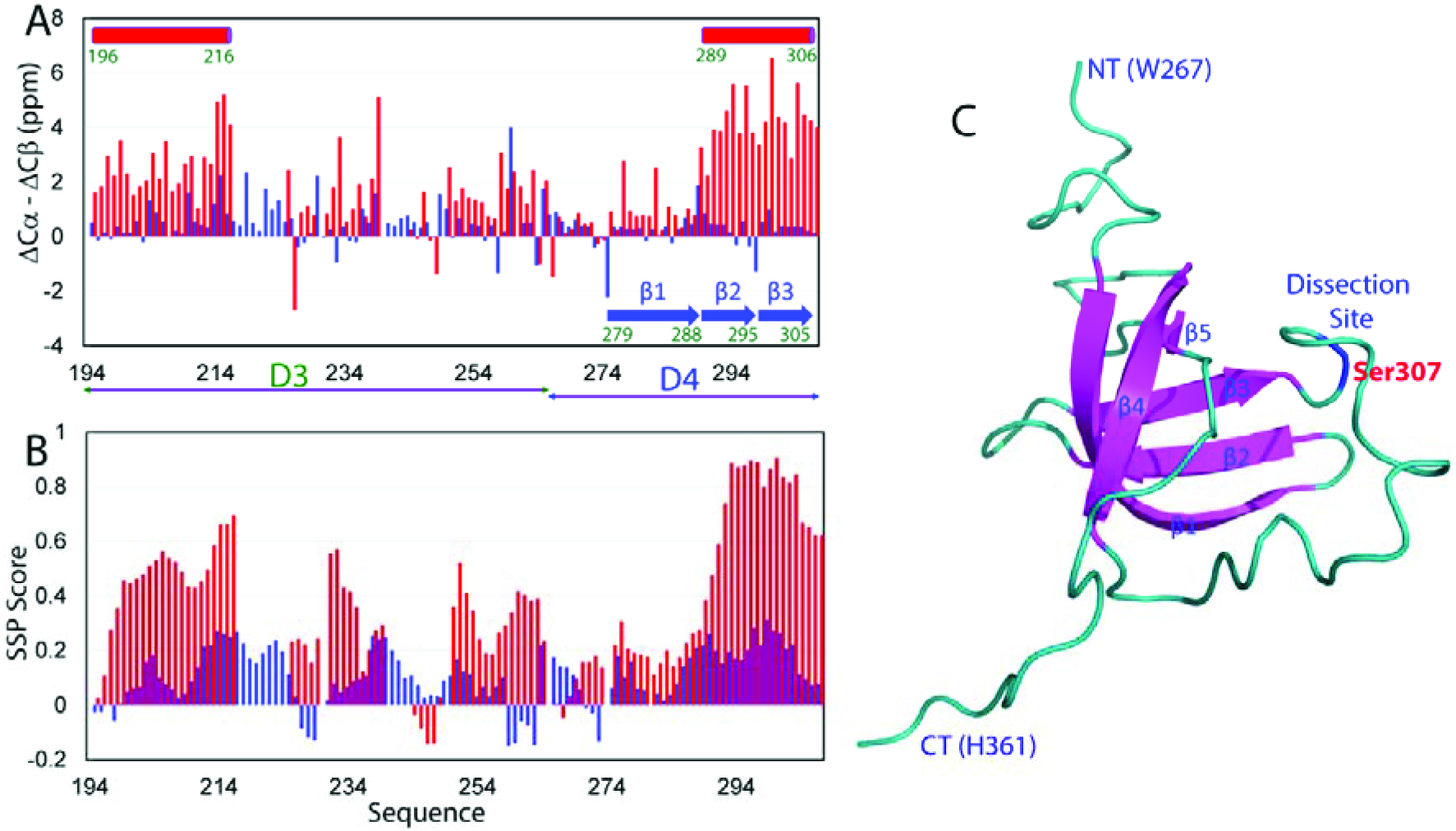
Residue-specific conformations of F4. (A) Residue specific (ΔCα-ΔCβ) chemical shifts of F4 in in aqueous solution (blue) and in DMPC/DHPC bicelle (red). (B) Secondary structure scores of F4 in in aqueous solution (blue) and in DMPC/DHPC bicelle (red), which were obtained by analyzing their chemical shifts with the SSP program. A score of + 1 is for the well-formed helix while a score of -1 for the well-formed extended strand. (C) NMR structure of the forth S1 domain with the dissection site labeled.

By a sharp contrast, upon interacting with bicelle, many residues of F4 suddenly had large and positive (ΔCα–ΔCβ) chemical shifts (Fig 6A), indicating that they became highly helical (Fig 6B). In particular, the N-terminal residues Val195-Val214 and C-terminal residues Tyr290-Val306 became highly helical as quantitatively reflected by their positive and large SSP scores (Fig 6B). The results together provide residue-specific evidence that upon interacting with bicelle, many residues of F4, particular the N- and C-terminal regions undergo a conformational transition from a predominantly disordered state to highly helical conformation.

### 4. Principle mediating the formation of membrane-embedded amphiphilic helices

To decode the principle mediating the formation of the helical conformations of the N- and C-terminal regions of F4 upon interacting with bicelle, we plotted their helical wheel diagrams for residues Val195-Val214 (Fig 7A) and Tyr290-Val306 (Fig 7B). Indeed, both of them are predicted to have capacity in adopting “amphiphilic helix”, which is characterized by spatial segregation of polar and non-polar amino acids that are located on opposing faces as oriented along the long axis of the helix. Amphiphilic helix was first identified in myoglobin and haemoglobin in 1965 [68], and its importance in mediating protein-lipid interactions was first recognized in 1974 [69]. Hydrophobic moment, or amphiphilicity, reflecting the periodicity in protein hydrophobicity, was proposed to detect the ability of a protein sequence to form amphiphilic helix [70,71], which has been now demonstrated to mediate various protein-membrane interactions [70-74]. Amazingly, bioinformatics studies revealed that all proteins universally have segments with high intrinsic amphiphilicity even including randomly generated sequences, regardless of their native structures [75,76].

Indeed, based on dihedral angle constraints derived from the NMR chemical shifts, we modeled the structures of the two fragments and both of them form amphiphilic helice (Fig 7C and 7F), with one side being largely non-polar (Fig 7D and 7G); and another side highly polar (Fig 7E and 7H).

**Fig 7.**
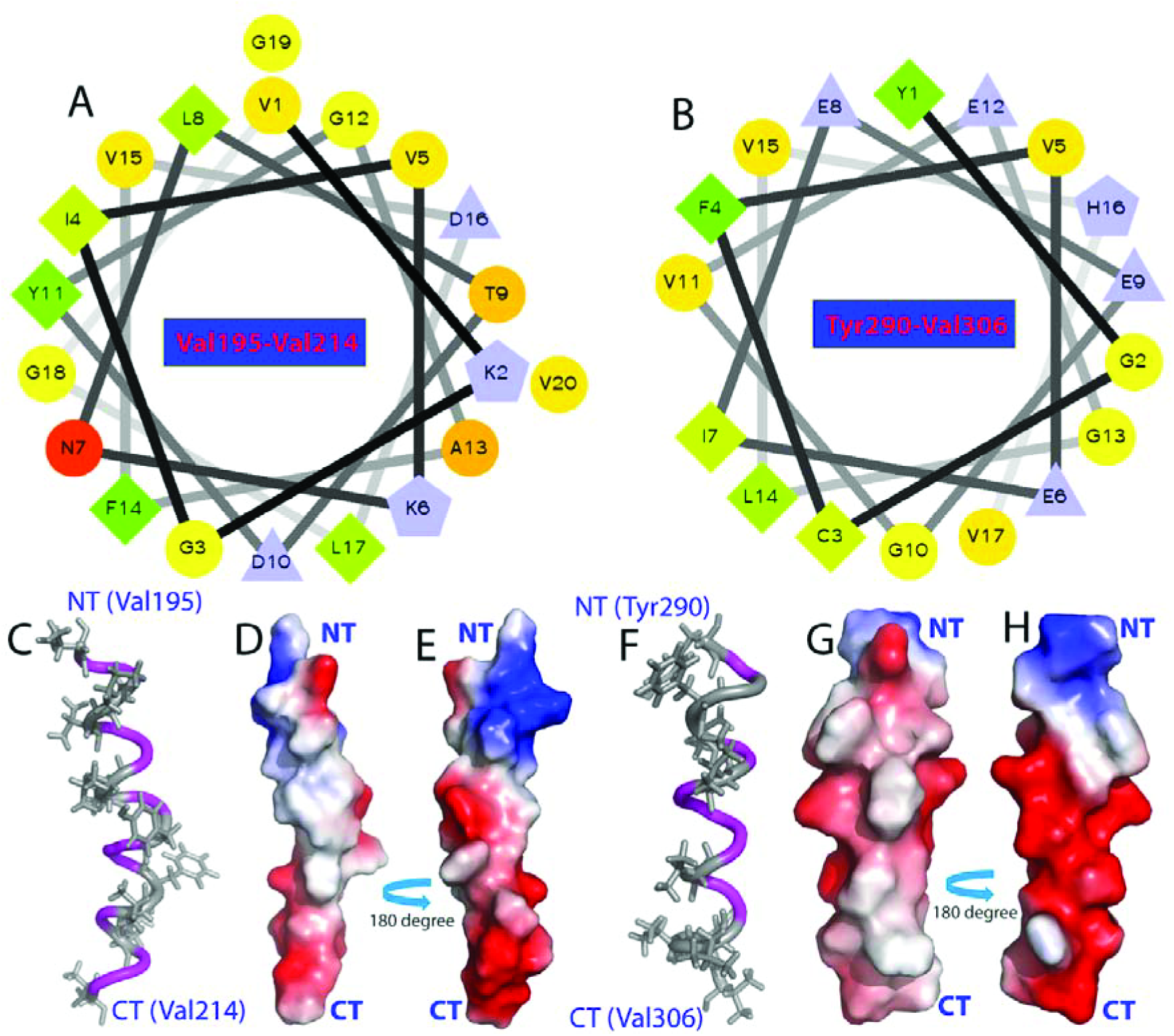
Visualization of two amphiphathic helices of F4 formed in bicelle. Helical wheel diagrams of the segments over residues Val195-Val214 (A) and Tyr290-Val360 (B); which are characterized by spatial segregation of polar and non-polar amino acids that are located on opposing faces as oriented along the long axis of the helix. Three-dimensional models and their electrostatic potential surfaces respectively for Val195-Val214 (C-E) and Tyr290-Val360 (F-H) in bicelle.

We calculated the hydrophobic moment or amphiphilicity of F4 and interestingly, even for such a short fragment, four regions have high amphiphilicity, which include Val195-Val214, His219-Leu246, Trp267-Thr285 and Tyr290-Val360 (Fig 8A), consistent with previous reports that amphiphilic fragments universally exist in all proteins [75,76]. However,why out of the 4 segments, only the N- and C-terminal segments were experimentally shown to have high capacity in interacting with bicelle ?

We thus calculated the hydrophobicity scale [77] of F4 (Fig 8B), and interestingly the middle two segments over His219-Leu246 and Trp267-Thr285 have negative hydrophobicity scales for most of their residues. Indeed, examination of the helical wheel diagrams (Fig 8C and 8D) reveals that their hydrophobic surfaces are much smaller than those of Val195-Val214 (Fig 7A) and Tyr290-Val306 (Fig 7A). This observation immediately suggests that both amphiphilicity and hydrophobicity need to be sufficiently high for a protein segment to have strong ability in forming a membrane-induced amphiphilic helix. To confirm this finding, we also calculated the amphiphilicity and hydrophobicity for the rest of the dissected fragments (Fig 9).Indeed, only F5 has two segments with both amphiphilicity and hydrophobicity being relatively large (Fig 9), which is completely consistent with the CD results that F5 also had large conformational changes upon interacting with bicelle (Fig 4).

**Fig 8.**
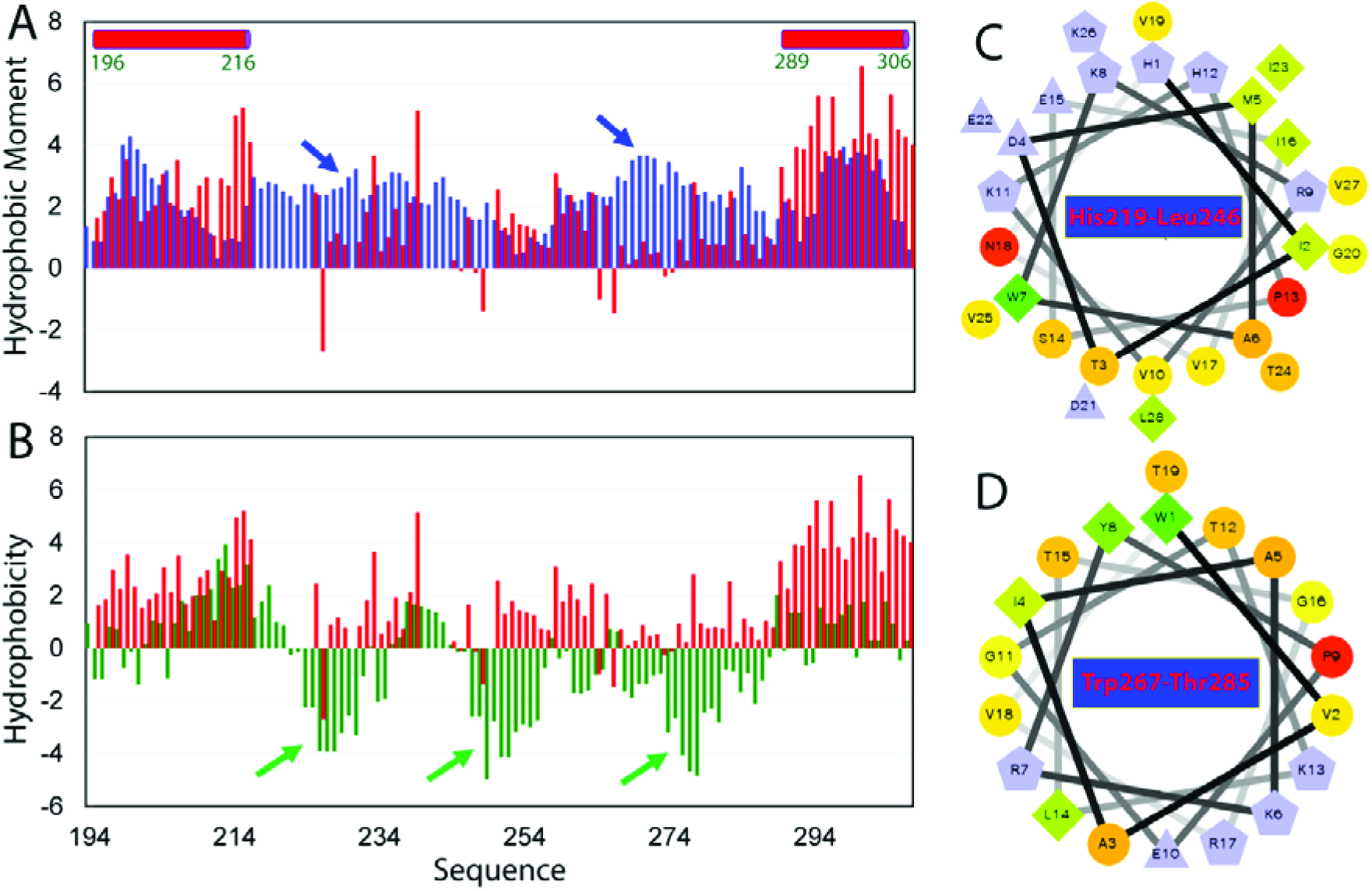
Membrane-interactions mediated by both amphiphilicity and hydrophobicity. Amphiphilicity (blue) and residue specific (ΔCα-ΔCβ) chemical shifts of F4 in DMPC/DHPC bicelle (red). (B) Hydrophobicity (blue) and residue specific (ΔCα-ΔCβ) chemical shifts of F4 in DMPC/DHPC bicelle (red). Helical wheel diagrams of the segments over residues His219-Leu246 (C) and Trp267-Thr285 (D). To make them comparable with the (ΔCα-ΔCβ) chemical shifts, the values of Amphiphilicity and Hydrophobicity times a factor of 3.

**Fig 9.**
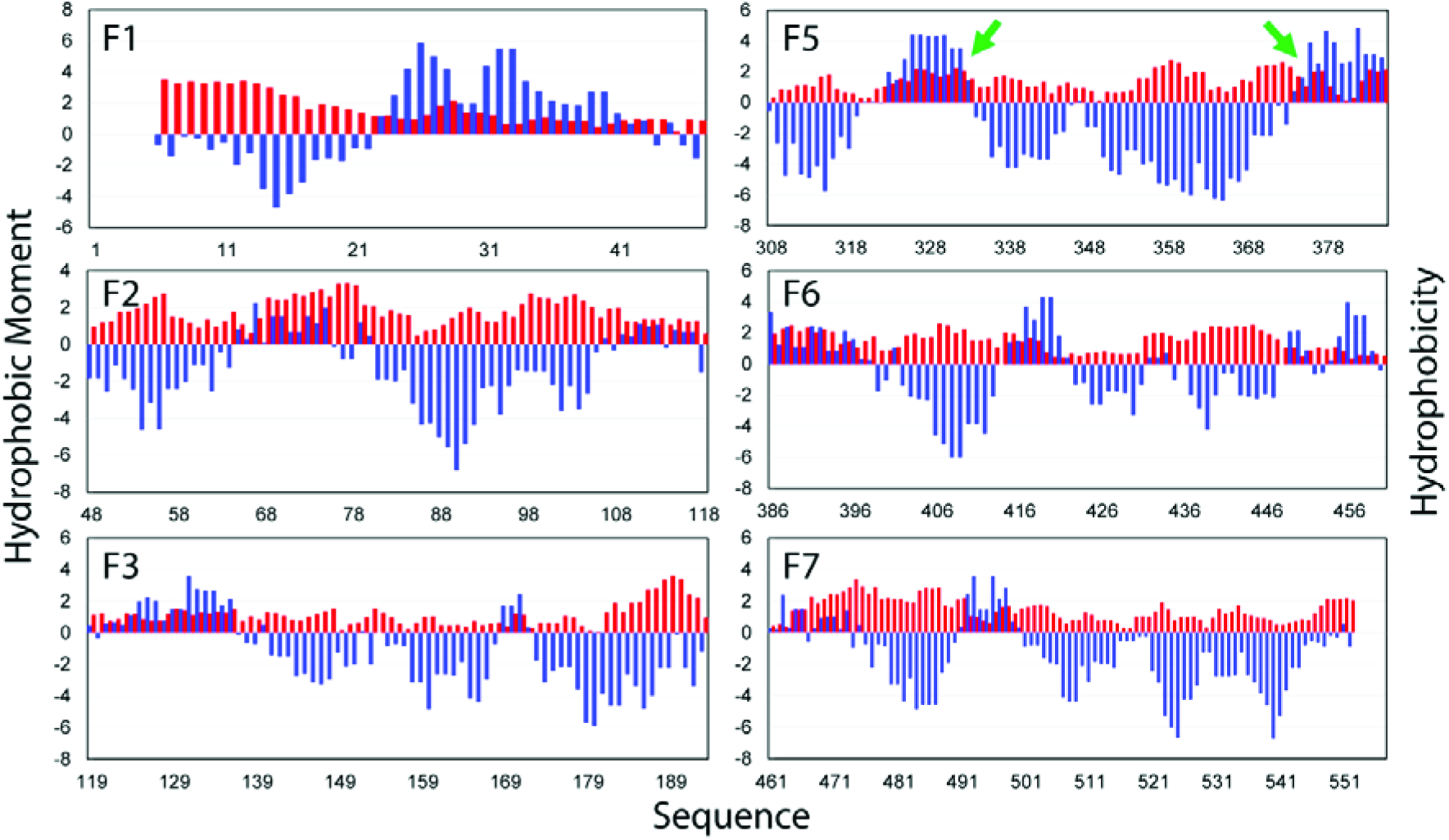
Amphiphilicity and Hydrophobicity of different S1 fragments. (A) Hydrophobicity (blue) and Amphiphilicity (red) of 6 dissected fragments of the *E. coli* S1 ribosomal protein. The values of Amphiphilicity and Hydrophobicity times a factor of 3.

**Fig 10.**
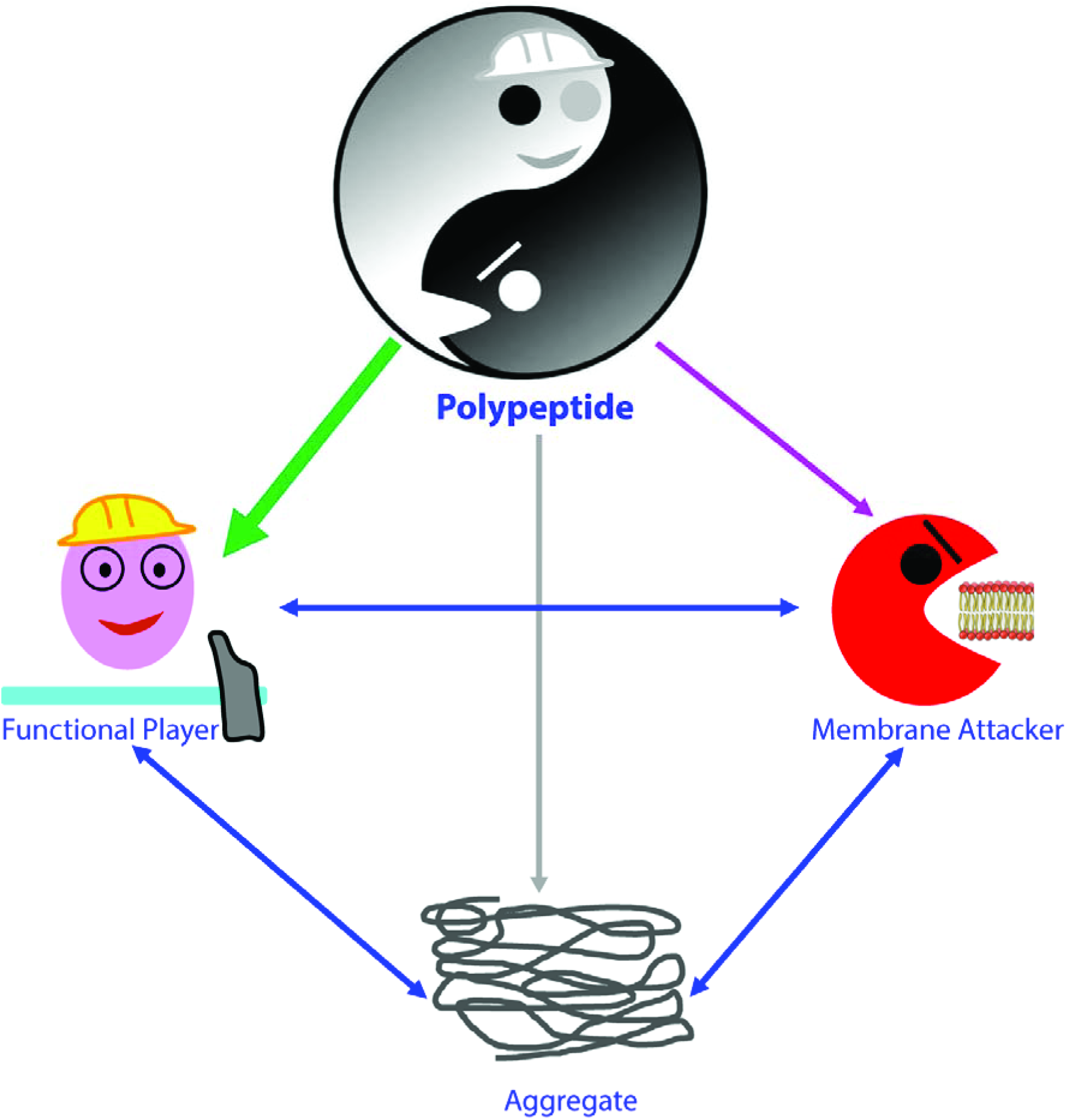
Proposed diagram for transformation of a protein from a functional player to an evil by unlocking the potential capacity to attack membranes, which is universally characteristic of severe aggregation.

However, considering the fact such diverse amphiphiphic regions are involved in protein-lipid interactions [73], it is possible that other fragments other than F4 are also interact with membranes/lipids with specific chemical properties.

## Discussion

There are around 300 hypotheses of aging and the best known is the free radical hypothesis. Proteins are highly susceptible to oxidative damages which lead to many irreversible covalent modifications including fragmentation of protein backbones [60-63]. In particular, during aging of *E. coli* cells, oxidative damage represents a major factor in triggering aggregation of many non-specific proteins [53-56,63]. Unfortunately, the mechanism underlying the oxidation-induced protein aggregation still remains highly elusive.

Here, to mimic the oxidation-induced protein fragmentation, we dissected the *E. coli* S1 ribosomal protein into 7 fragments. Detailed CD and NMR studies reveal that the dissection did eliminate all 6 well-structured S1 domains. However, out of 7 fragments, only F4 becomes “completely insoluble” while the other 6 fragments are still soluble in buffer to some extent. As all 7 fragments contain parts of the well-folded regions in 6 all-β S1 domains, this implies that despite the elimination of the tertiary folds, fragments except for F4 can rearrange their secondary structures from β-strands into helices to bury hydrophobic regions in order to avoid “complete insolubility”, as we previously demonstrated on a Nck2 SH3 domain [83]. On the other hand, because F4 has longer regions which have both high amphiphilicity and hydrophobicity (Fig 8 and 9), it thus appears impossible for F4 to have them all properly buried. As a consequence, it becomes “completely insoluble” even in 1 mM phosphate buffer. Furthermore, we observed that other fragments also started to precipitate in buffers containing high concentrations of salts. Therefore, our current results with protein fragmentation enforce our previous discovery that modifications such as a point mutation or cofactor depletion in β-rich proteins such as S1, SH3, MSP and SOD1 folds, is sufficient to completely eliminate their tertiary structures, which accounts for reduction of solubility or even becoming “completely-insoluble” [31-52]. This mechanism also rationalizes previous observations that β-rich proteins were significantly over-represented in aggregation-prone proteins relative to the proteomes [81], because the tertiary folds rich in β-sheets are relatively easier to be abolished by sequence modifications [31-52].

Remarkably, the asymmetric segregation of protein aggregates in *E. coli* led to the aged mother cells holding the accumulated aggregates and rejuvenated daughter cells free of aggregates [17]. This implies that the aggregation-prone proteins may also trigger cellular aging of *E. coli* cells by a “gain of cytotoxicity” mechanism. So a fundamental question remaining to be answered is: whether the elimination of the well-folded structures of *E. coli* cytosolic proteins will also lead to gain of a novel capacity to abnormally interact with membrane? Here we experimentally reveal that the fragments like F4 resulting from the oxidation-induced fragmentation of an *E. coli* cytosolic protein do acquire a novel and strong capacity in abnormally interacting with membranes, if the fragments have both high amphiphilicity and hydrophobicity, which are thus prone to severe aggregation in cells or buffers [31-52]. In other words, the proteins/fragments characteristic of severe aggregation also own high capacity to attack membranes, as we previously decoded for ALS-causing mutants [32,43-45,50,52]. Consequently, this suggests that to abnormally interact with membranes may also represent a common mechanism by which aggregation-prone proteins cause aging by a “gain of cytotoxicity” mechanism.

Our current study on a protein of *E. coli*, the lowest-level organism, decrypt that the *E. coli* protein also follows the same mechanism of aggregation and gain of membrane-toxicity as the proteins of human being, the highest organism [31-52]. Therefore. proteins, the most important functional players for all forms of life, have potentials to transform from functional players to evils triggering diseases and aging by abnormally attacking membranes, if their hydrophobic/amphiphilic regions are unlocked by genetic, pathological or/and environmental factors, which is characteristic of severe aggregation. So the proteome appears to be Pandora’s box.

It has been shown that in *E. coli*, most cytosolic proteins need to fold into welldefined structures for their functions [84-85], while in eukaryotic organisms there are intrinsically disordered proteins [5], or even intrinsically insoluble proteins [32], which were sent for degradation upon synthesis [86-87]. In cells, intrinsically-insoluble, misfolded, and structure-eliminated proteins are expected to rapidly form aggregates due to a high salt concentration (∼150 mM) in cellular environments [31-52]. However, although the chance might be extremely rare, if some of these aggregation-prone proteins/fragments escape from forming “completely insoluble” aggregates, they may transform into evils to trigger diseases and aging by attacking membranes. This also explains the recent observation that human diseases were initiated by the soluble oligomers but not the insoluble aggregates or amyloid fibers, as the formation of soluble oligomers may facilitate these proteins to access membranes. Although it has been recently revealed that the abnormal insertion of SOD1 mutants into ER membranes without forming detectable aggregates is sufficient to initiate ALS pathogenesis [18], the coupling of the membrane-interaction and aggregation/amyloid-formation onto/within membranes may significantly strengthen the capacity of aggregation-prone proteins to cause diseases and aging [45,50,52], in particular for *E. coli* cells which do not have a complex membrane-network within their cellular space.

## Methods

## Generation of Recombinant Proteins

The DNA fragment encoding *E. coli* S1 ribosomal protein was amplified by PCR reaction directly on *Escherichia coli* BL21 (DE3) cells (Novagen) with designed primers and subsequently cloned into a modified vector pET28a with 6 His residues at C-terminus as we extensively used for the TDP-43 prion-like domain [51]. Furthermore, DNA fragments encoding 7 dissected fragments (Fig 1) were successfully obtained by PCR reactions with designed primers and subcloned into the same vector. The expression vectors were subsequently transformed into and overexpressed in *Escherichia coli* BL21 (DE3) cells (Novagen). The recombinant proteins were found inclusion body, and consequently were purified by a Ni^2+^-affinity column (Novagen) under denaturing conditions in the presence of 8 M urea. The fractions containing the recombinant proteins were acidified by adding 10% acetic acid and subsequently purified by reverse-phase (RP) HPLC on a C4 column eluted by water-acetonitrile solvent system. The HPLC elutions containing pure recombinant proteins were lyophilized and stored in -80 °C.

The generation of the isotope-labelled proteins for NMR studies followed a similar procedure except that the bacteria were grown in M9 medium with the addition of (^15^NH_4_)_2_SO_4_ for ^15^N labeling and (^15^NH_4_)_2_SO_4_/[^13^C]-glucose for double labelling [31-52]. The purity of the recombinant proteins was checked by SDS–PAGE gels and their molecular weights were verified by a Voyager STR matrix-assisted laser desorption ionization time-offlight-mass spectrometer (Applied Biosystems). The concentration of protein samples was determined by the UV spectroscopic method in the presence of 8 M urea. Briefly, under the denaturing condition, the extinct coefficient at 280 nm of a protein can be calculated by adding up the contribution of Trp, Tyr and Cys residues [88].

## CD and NMR experiments

All circular dichroism (CD) experiments were performed on a Jasco J-810 spectropolarimeter equipped with a thermal controller using 1-mm path length cuvettes. Data from five independent scans were added and averaged. CD samples were prepared by diluting the concentrated samples dissolved in Milli-Q water (pH 4.0) into 1 mM phosphate buffer to reach the final concentration of 15 μM at pH 6.8.

To assess the capacity of the full-length S1 protein and its 7 dissected fragments to interact with membranes, here we used DMPC/DHPC bicelle to mimic the bilayer membrane, which was prepared by mixing up dimyristoylphosphatidylcholine (DMPC) and dihexanoylphophatidylcholine (DHPC) at a q value of 0.25 as previously described [43,51]. The liposome was prepared as we previously described [43] but with the total extract of *E. coli* lipids.

All NMR experiments were acquired on an 800 MHz Bruker Avance spectrometer equipped with pulse field gradient units as described previously [31-52,89,90]. For characterizing the residue-specific conformations of the fragment 4 in both aqueous solutions and bicelle, two pair of triple-resonance experiments HN(CO)CACB, CBCA(CO)NH were collected for the sequential assignment on ^15^N-/^13^C-double labelled samples respectively in Milli-Q water at pH 4.0 and in bicelle at protein concentration of 300 μM. NMR data were processed with NMRPipe [91] and analyzed with NMRView [92].

## Structure Modelling

To visualize the conformation in bicelle of two helical segments of F4 respectively over residues Val195-Val214 and Tyr290-Val360, the backbone dihedral angles were generated with TALOS+ by inputting backbone ^1^H, ^15^N and ^13^C chemical shifts [93], which were utilized to generate the model of two segments by CYANA [45,48,50,52,94]. The lowest target-function CYANA structures with no dihedral angle violation >4 degrees were selected for analysis.

## Author Contributions

Conceived and designed the experiments: JXS; Performed the experiments: LZL YML. Analyzed the data: JXS LZL YML. Prepared figures and wrote the paper: JXS.

## Acknowledgement

This study is supported by Ministry of Education of Singapore (MOE) Tier 2 Grants 2011-T2-1-096 and MOE2015-T2-1-111 to Jianxing Song. The funders had no role in study design, data collection and analysis, decision to publish, or preparation of the manuscript.

